# TweetyBERT: Automated parsing of birdsong through self-supervised machine learning

**DOI:** 10.1101/2025.04.09.648029

**Authors:** George Vengrovski, Miranda R. Hulsey-Vincent, Melissa A. Bemrose, Timothy J. Gardner

## Abstract

Deep neural networks can be trained to parse animal vocalizations – serving to identify the units of communication, and annotating sequences of vocalizations for subsequent statistical analysis. However, current methods rely on human labelled data for training. The challenge of parsing animal vocalizations in a fully unsupervised manner remains an open problem. Addressing this challenge, we introduce TweetyBERT, a self-supervised transformer neural network developed for analysis of birdsong. The model is trained to predict masked or hidden fragments of audio, but is not exposed to human supervision or labels. Applied to canary song, TweetyBERT autonomously learns the behavioral units of song such as notes, syllables, and phrases - capturing intricate acoustic and temporal patterns. This approach of developing self-supervised models specifically tailored to animal communication will significantly accelerate the analysis of unlabeled vocal data.

## Introduction

Artificial intelligence offers great promise to enhance our understanding of animal vocalizations by enabling analysis of behavioral recordings at unprecedented scales (Rutz et al., 2023). Machine learning algorithms have already demonstrated substantial progress in classifying species-specific vocalizations, with applications such as Merlin Bird ID enabling species identification from environmental recordings (Kahl et al., 2021). However, the current challenge lies in developing methods that go beyond species identification to parse animal vocalizations into their fundamental behavioral units—notes, syllables, motifs, and bouts—that collectively structure complex vocal sequences (Cohen et al., 2022; Gu et al., 2023; Sainburg et al., 2020). Segmentation of vocalizations into these discrete behavioral units will enable detailed modeling of communication signals, linking specific vocal elements to behavioral contexts, emotional conditions (Briefer et al., 2022), or environmental stimuli. For basic neuroscience, achieving this level of analysis can help decode the “syntax” and “grammar” of animal communication, allowing for more detailed studies of the neural and cognitive mechanisms underlying vocal behavior (Cohen et al., 2020; Kobayashi et al., 2001; Koparkar et al., 2024; Markowitz et al., 2013; Veit et al., 2020).

Supervised and semi-supervised methods for song labelling have been pursued for many years (Daou et al., 2012; Koumura & Okanoya, 2016; Tachibana et al., 2014). Current deep neural networks are trained end-to-end to produce labels based on raw inputs, but these networks require human-labeled datasets for training (Cohen et al., 2022; Steinfath et al., 2021). This reliance on manual annotations severely restricts scalability and generalizability. To circumvent these limitations, unsupervised methods seek to develop song representations directly from acoustic data without human annotation (Goffinet et al., 2021; Sainburg et al., 2020; Singh Alvarado et al., 2021). An essential step in this unsupervised parsing of animal vocal sequences involves creating a latent feature space (or embedding) in which acoustically similar sounds cluster together. However, existing unsupervised approaches, such as variational autoencoders (VAE) and direct dimensionality reduction techniques like Uniform Manifold Approximation and Projection (UMAP; McInnes, Healy, and Melville, 2018) have a limited capacity to deal with the real-world variability of natural vocalizations. Existing algorithms typically start by splitting song into artificial units that are roughly the duration of a single syllable or vocal utterance. Similarity scores are computed between these fragments and form the basis of latent space analysis or clustering. This approach of fragmenting song into small blocks of a fixed duration works for some of the simplest or most stereotyped animal vocalizations, but struggles to identify vocal clusters with significant temporal and acoustic variations (Coffey et al., 2019; Goffinet et al., 2021; Kollmorgen et al., 2020; Sainburg et al., 2020).

In contrast, transformer models originally developed for natural language processing excel at capturing complex temporal relationships among elements within sequences (Vaswani et al., 2017). Rather than imposing fixed windows, transformers use self-attention mechanisms and positional encodings to robustly model long-range contextual relationships, a property that has revolutionized self-supervised learning in language (Devlin et al., 2018; Radford & Narasimhan, 2018; Touvron et al., 2023). These architectures have successfully generalized beyond text-based tasks, and are commonly applied to speech, vision, and multimodal information, as demonstrated by models like wav2vec2.0 and HuBERT (Baevski et al., 2020; Dosovitskiy et al., 2020; Hsu et al., 2021). Applying transformer-based methods to animal communication could overcome limitations posed by prior fixed-window methods and better accommodate natural variability in animal vocalizations. However, existing transformer models applied to bioacoustics tasks usually require supervised fine-tuning on human-labeled data, thus inheriting the practical constraints of other supervised methods (Gu et al., 2023; Hagiwara, 2023).

Here, we introduce TweetyBERT, a transformer-based neural network specifically designed to process the fast, complex vocalizations of songbirds. The model operates at a 10x higher temporal resolution than most human speech models but is trained with a similar masked prediction task - predicting held-out fragments of the audio recording. Focusing on canary song (Fig. 1) we demonstrate that TweetyBERT spontaneously learns representations of individual syllables as distinct elements within its latent embedding space. Remarkably, this structured representation emerges solely from a masked prediction task, without explicit supervision on syllable categories or boundaries and without additional fine-tuning. These embeddings facilitate automated clustering of vocal elements that closely align with human annotations. Furthermore, TweetyBERT’s latent space exhibits high consistency for individual canaries recorded at different times within the spring breeding season, but also reveals significant variations between breeding season song and fall plastic song (Nottebohm et al., 1986; Voigt & Leitner, 2008). Whether through automated clustering of vocalizations, or qualitative analysis of latent space trajectories, self-supervised transformer models developed specifically for animal vocalizations will open new doors in the analysis of animal communication.

**Fig. 1.**
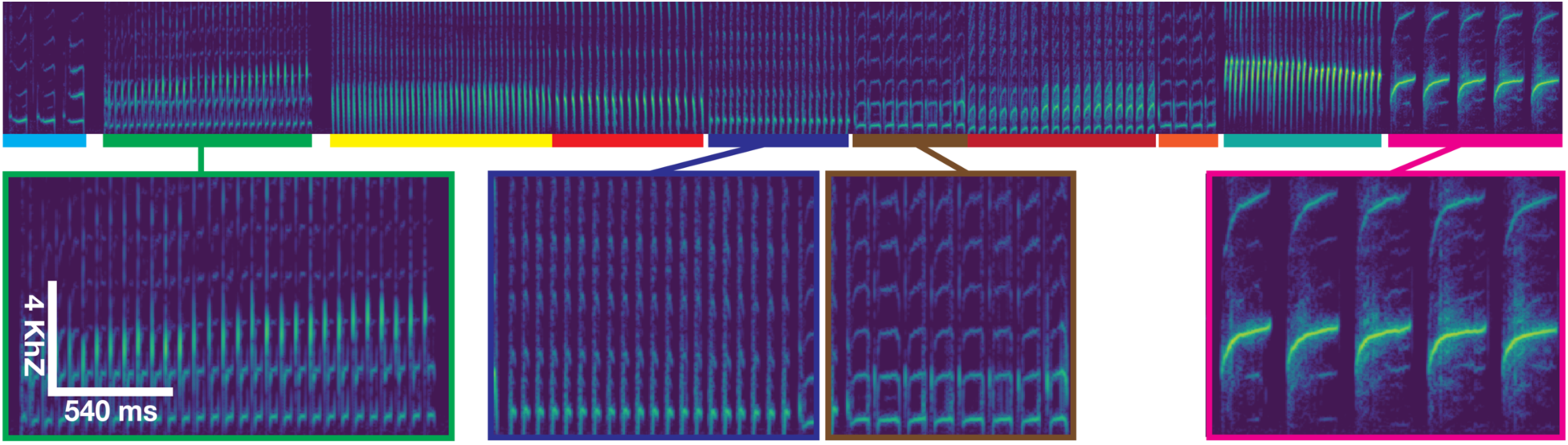
Typical canary song in spectrogram form. Canary song consists of stuttered syllables grouped in phrases (color bars) arranged sequentially.

## Results

### Model Architecture

TweetyBERT is a compact (2.5 million parameter), spectrogram-based, self-supervised transformer network inspired by the architectures of BERT and TERA (Devlin et al., 2018; Liu et al., 2020). Unlike many of the large speech transformers (Baevski et al., 2020; Hsu et al., 2021), TweetyBERT operates directly on spectrograms without any discretization or vector quantization, simplifying the pre-training process and reducing the number of hyperparameters influencing network performance. It integrates a lightweight convolutional front-end designed to extract spectral features, as convolutional layers effectively model local acoustic patterns and improve transformer-based architectures (Gulati et al., 2020). This convolutional front-end preserves temporal dimensionality, facilitating a direct alignment between internal representations and corresponding input spectrogram frames for subsequent analysis. The model’s output dimensions precisely match the input spectrogram, facilitating pixel-level reconstruction. The training objective is to minimize the mean squared error between predicted and actual pixel values within masked regions of the spectrogram.

### Input Representation and Training Procedure

For training data, we utilized audio recordings from three canaries previously described by (Cohen et al., 2022). A single TweetyBERT model was trained jointly on data from all three birds, comprising approximately 15 hours of song (Supplementary Fig. 1). Song segments were isolated from these recordings using a supervised ‘song detector’ (Supplementary Fig. 2). This song detector eliminated calls, cage noise, and extended silences typically present in raw recordings. These songs were converted to spectrograms with a 2.7 ms time resolution (the time width of a single column of the spectrogram image). Throughout this paper, the terms “frames” and “points” all refer to these discrete time bins. During training, we apply a variable masking strategy, randomly masking 25% of each spectrogram segment to encourage learning at multiple temporal scales (Supplementary Fig. 3). To further improve robustness and encourage translational invariance, we augmented the training dataset by applying random frequency shifts and extracting multiple random 2.7 second temporal crops from each audio segment. Consequently, the model encountered varied representations of every song segment during training, enhancing its ability to generalize across temporal positions, or frequency variability.

### Analysis and Representation Extraction

TweetyBERT’s masked-prediction training is intended to encourage the network to form meaningful internal representations of canary songs; the reconstruction task itself is discarded after training, as is typical in self-supervised learning. Our primary insights come from analyzing the model’s internal representations, not its output predictions (Fig. 2, Inference phase). Internal representations are extracted as activation vectors corresponding to intermediate model layers. To analyze longer sequences or multiple songs, we concatenate these activation vectors. For visualization and further analysis, we project these concatenated high-dimensional vectors into two dimensions using UMAP dimensionality reduction.

**Fig. 2.**
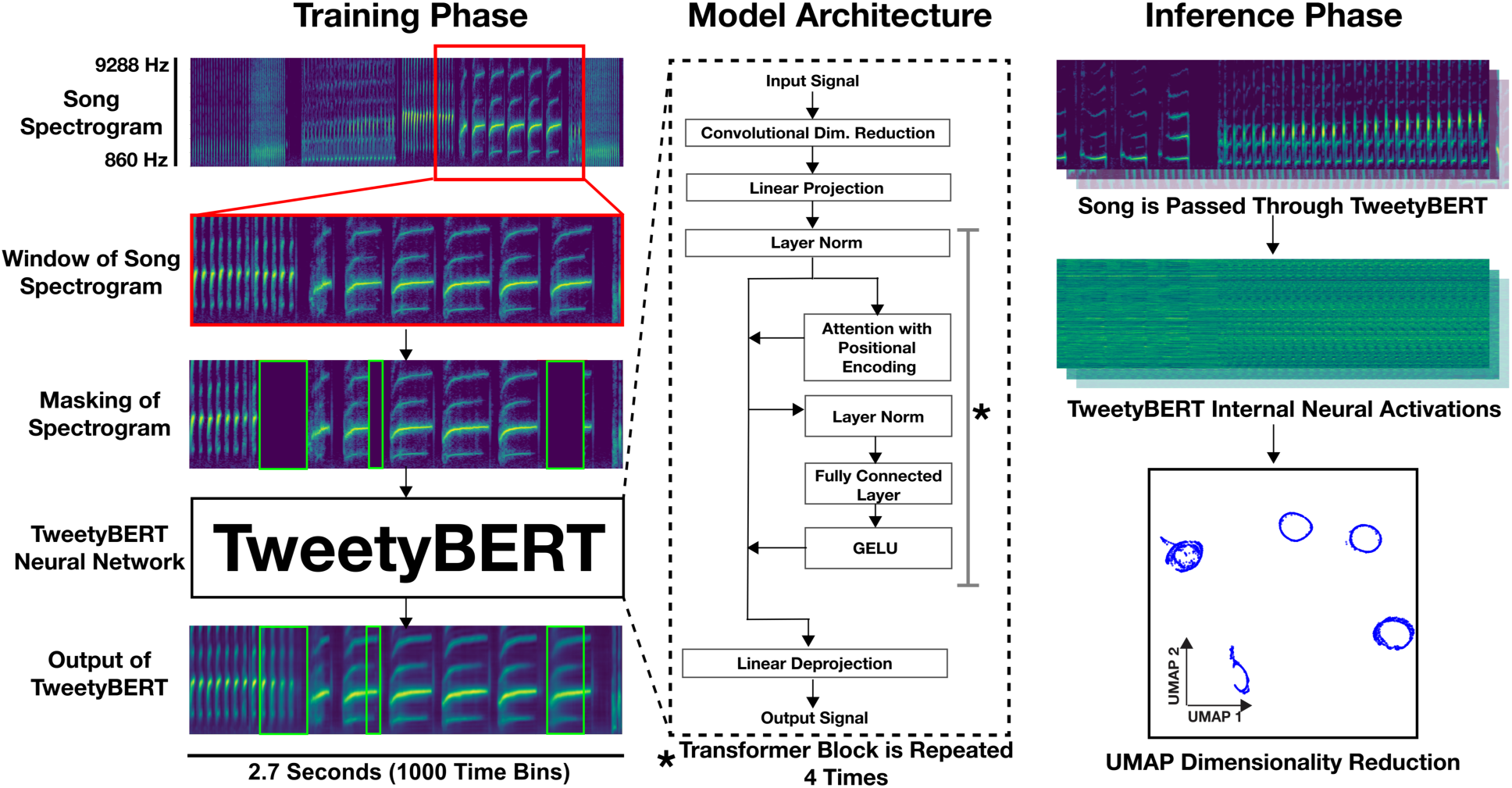
TweetyBERT architecture. **Training phase:** Random segments (2.7 s) of canary song spectrograms are masked and input to the TweetyBERT network, which learns to reconstruct masked spectrogram regions. **Model architecture:** TweetyBERT is a self-supervised transformer network operating directly on spectrogram inputs, combining a convolutional front-end (for local acoustic feature extraction) with a compact transformer backbone. **Inference phase:** Song spectrograms are passed through the trained network, and intermediate neural activations are extracted for analysis. The UMAP visualization shows that activations corresponding to distinct syllable types typically form elliptical trajectories in reduced dimensional space.

### Emergence of Structured Representations in TweetyBERT Embeddings

If birdsong syllables consisted of simple notes such as the sounds produced by piano keys, then dimensionality reduction methods (such as UMAP) applied directly to spectrogram images would clearly separate syllables into distinct clusters. However, canary syllables—and animal vocalizations more generally—require a consideration of the time-course of acoustic elements to properly categorize vocalizations. This complexity is illustrated in (Fig. 3C), where UMAP dimensionality reduction was applied to spectrograms from three different canary songs. Here, each data point corresponds to a single spectrogram time bin, colored according to ground truth syllable labels from a previously described public dataset (Cohen et al., 2022). Black points are inter-syllabic silences. Different canary syllables share highly similar spectral features when analyzed at the single time bin level (Fig. 3C). In contrast, dimensionality reduction applied to TweetyBERT’s internal neural activations reveals a structured embedding space in which the units of song – syllables – are distinctly separable from one another (Fig. 3B). In this representation, syllables correspond to elliptical trajectories in the two-dimensional UMAP space. Points along an ellipse represent sequential temporal positions within a syllable’s acoustic trajectory, and a full traversal corresponds to the complete utterance of that syllable. Repeated syllables within a phrase trace the ellipse multiple times. Coincidentally, biophysical models of canary song found that most syllables could be produced by an elliptical trajectory in the space of air pressure and muscle tension (Gardner et al., 2001; Mindlin et al., 2003). The self-supervised transformer model appears to have found the simplest fundamental representation of canary syllables.

**Fig. 3.**
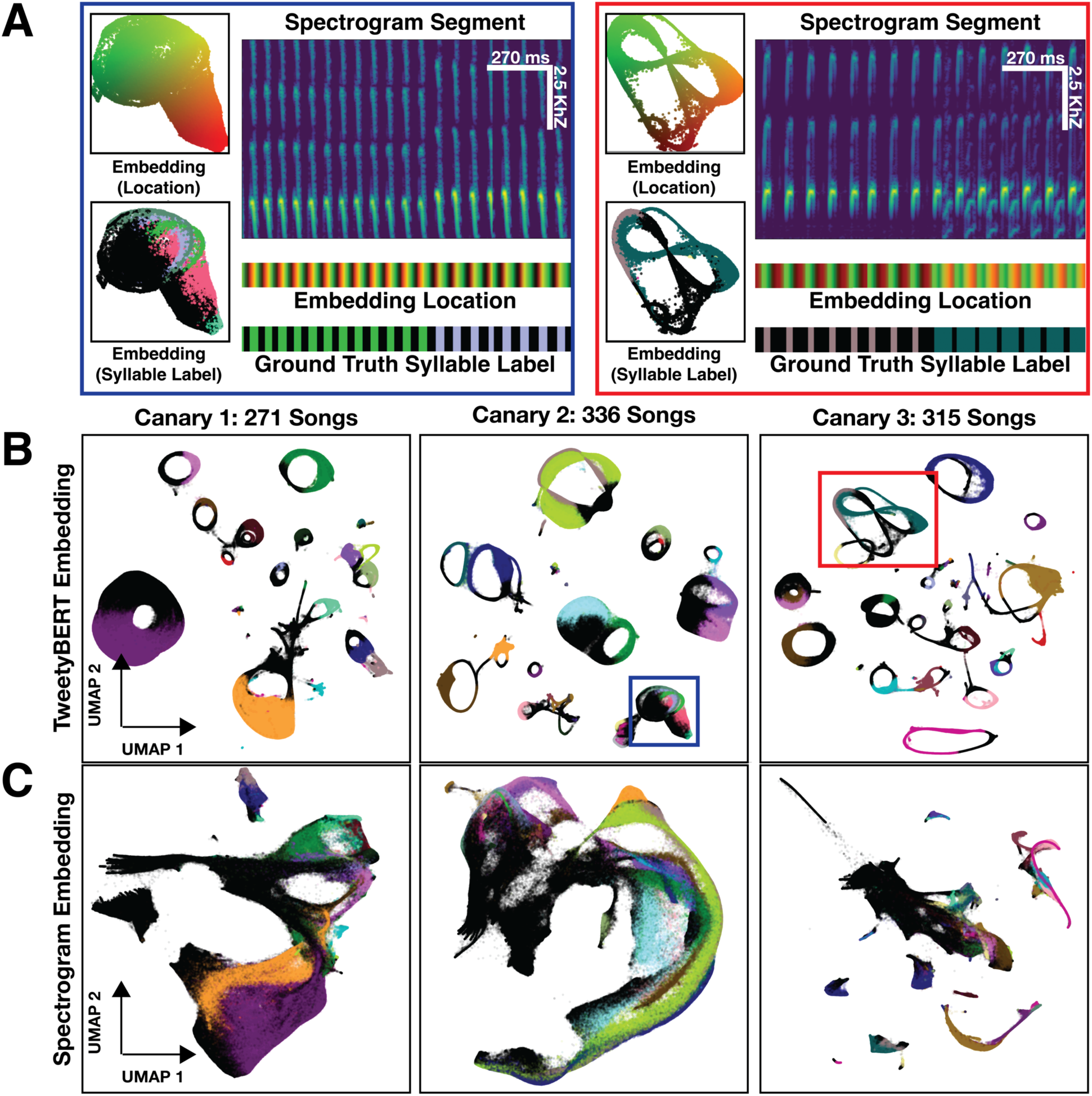
TweetyBERT and Spectrogram UMAP Embeddings. **(A)** Detailed analysis of syllables with more complex or overlapping latent space representations. In different panels, colors represent position in the latent space, or human ground truth labels. **(B)** Learned embeddings from three different canaries colored by human-annotated syllable classes; 1 million time bins. **(C)** Direct UMAP of spectrograms (baseline comparison); 1 million time bins.

### Acoustic Interpretation of Overlapping Ellipses

We occasionally observed regions of intersection or proximity between ellipses representing different syllable classes. Spectrogram analysis reveals that these overlapping regions often result from genuine acoustic similarities rather than limitations in the model’s representational capacity. For example, distinct syllable types sometimes share similar spectral properties during their onset, diverging only later in the syllable (Fig. 3A). This partial acoustic overlap leads to intersections in the embedding space, reflecting real acoustic relationships.

### Context-Dependent Representation of Silences

TweetyBERT encodes silences differently depending on their context within the song. Inter-syllabic silences—brief pauses between syllables—are embedded within the elliptical trajectories of their corresponding syllable classes. In contrast, extended silences found at song boundaries or between phrase transitions form a distinct, separate cluster that does not exhibit elliptical structure. Occasionally, this cluster also includes non-song elements, such as calls or background noise that were not fully excluded by the song detection step. This differentiation demonstrates the model’s capacity to distinguish silences that form part of a repeated syllable from longer periods of non-vocalization or noise.

### Clustering Methodology

TweetyBERT’s embedding space demonstrates a structured organization suitable for efficient clustering. We used Hierarchical Density-Based Spatial Clustering with Noise (HDBSCAN) due to the algorithm’s ability to automatically determine the number of clusters, adapt to variable cluster shapes, and handle noise effectively (Campello et al., 2013). Its computational complexity of *O*(*n log n*) makes it suitable for analyzing large datasets (up to ∼1 million time bins with 128 GB of RAM).

### Performance Evaluation

To quantitatively evaluate clustering performance, we applied the V-measure statistic (Rosenberg & Hirschberg, 2007), a metric previously used in zebra finch and Bengalese finch vocalization studies (Koch et al., 2024; Sainburg et al., 2020). V-measure is the harmonic mean of homogeneity (clusters contain one class) and completeness (each class forms a single cluster), ranging from 0 (no agreement) to 1 (perfect clustering). Clustering of TweetyBERT’s latent representations for three canaries (partitioned into 12 folds due to UMAP memory constraints) resulted in a high V-measure score of 0.88 ± 0.02. This data was drawn from held-out audio data not seen during training. The V-measure score confirms strong alignment between automated clustering and ground truth syllable classes, establishing a quantitative benchmark for unsupervised canary song clustering.

### Reconciling Labeling Schemes

A minor technical issue arose because our available ground truth labels existed at the syllable level, whereas HDBSCAN tended to cluster vocalizations at the phrase level. In canary song, a “phrase” is a sequence consisting of repeated renditions of the same syllable type. This discrepancy occurred primarily because HDBSCAN grouped inter-syllabic silences with their adjacent syllables, effectively merging syllables and their silences into larger, phrase-level clusters. To accurately evaluate clustering performance, we converted ground truth syllable labels into corresponding phrase-level labels as detailed in (Supplementary Fig. 4).

### Differences from Human Annotations

Although our clustering analysis showed strong overall agreement with human annotations, we identified specific common discrepancies. One difference was that HDBSCAN occasionally merged acoustically similar but distinct phrases into a single cluster. In many cases merged syllables are acoustically very similar and are challenging for expert human labelers to classify consistently. Occasionally, syllables appeared to be merged due to limitations of clustering in two-dimensional embedding space, where overlapping elliptical representations of distinct syllables were difficult to separate. Examples of these clustering differences are shown in (Fig. 4). One limitation of the current approach is that HDBSCAN clusters static UMAP embeddings without explicitly modeling their temporal structure, in contrast to dynamic methods such as Hidden Markov Models. Incorporating temporal dynamics explicitly into the clustering process could further reduce these discrepancies.

**Fig. 4.**
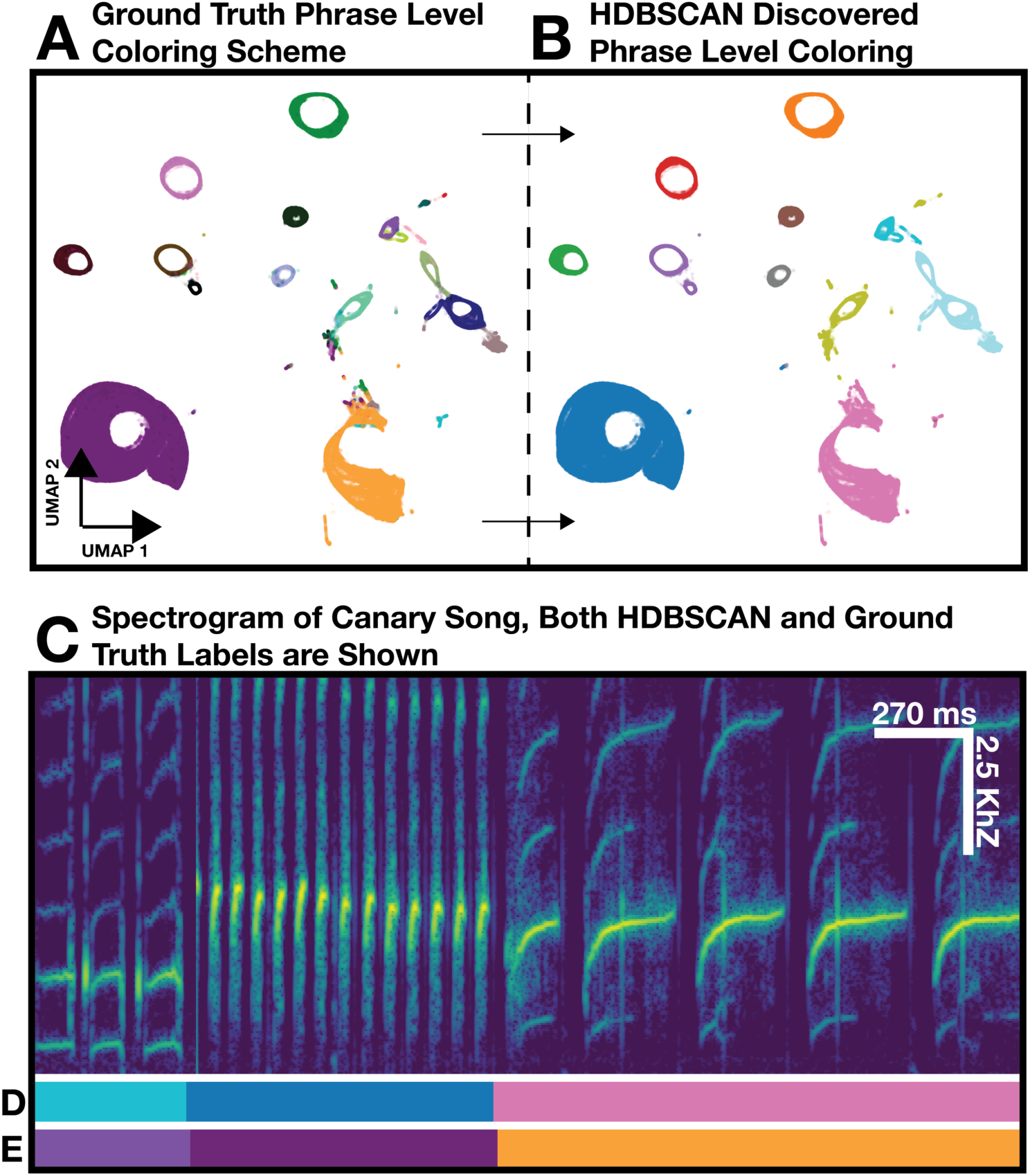
Machine derived clusters and human derived clusters are highly similar. **(A)** UMAP projection of TweetyBERT embeddings colored by human ground truth phrase labels. **(B)** Same embeddings colored by phrase clusters identified via HDBSCAN. **(C)** Spectrogram of canary song. **(D)** HDBSCAN-derived labels and **(E)** ground truth labels illustrating agreement between unsupervised clustering and manual annotations.

Finally, we also observed fragmentation of single phrases into multiple clusters, particularly in complex syllables. In these instances, the fragmentation accurately identified meaningful sub-units of long duration multi-part syllables (Fig. 5A & B). This fragmentation is arguably a more meaningful description of the long-multi-part syllables – and simply means that in these cases the transformer discovers a different effective “convention” for parsing song than the convention followed by the human labeler. Whether errors or different conventions, all these departures from human labels will lower the V-measure score. The match to human labels may also be further optimized by clustering in higher dimension, rather than projecting to 2D prior to clustering.

**Fig. 5.**
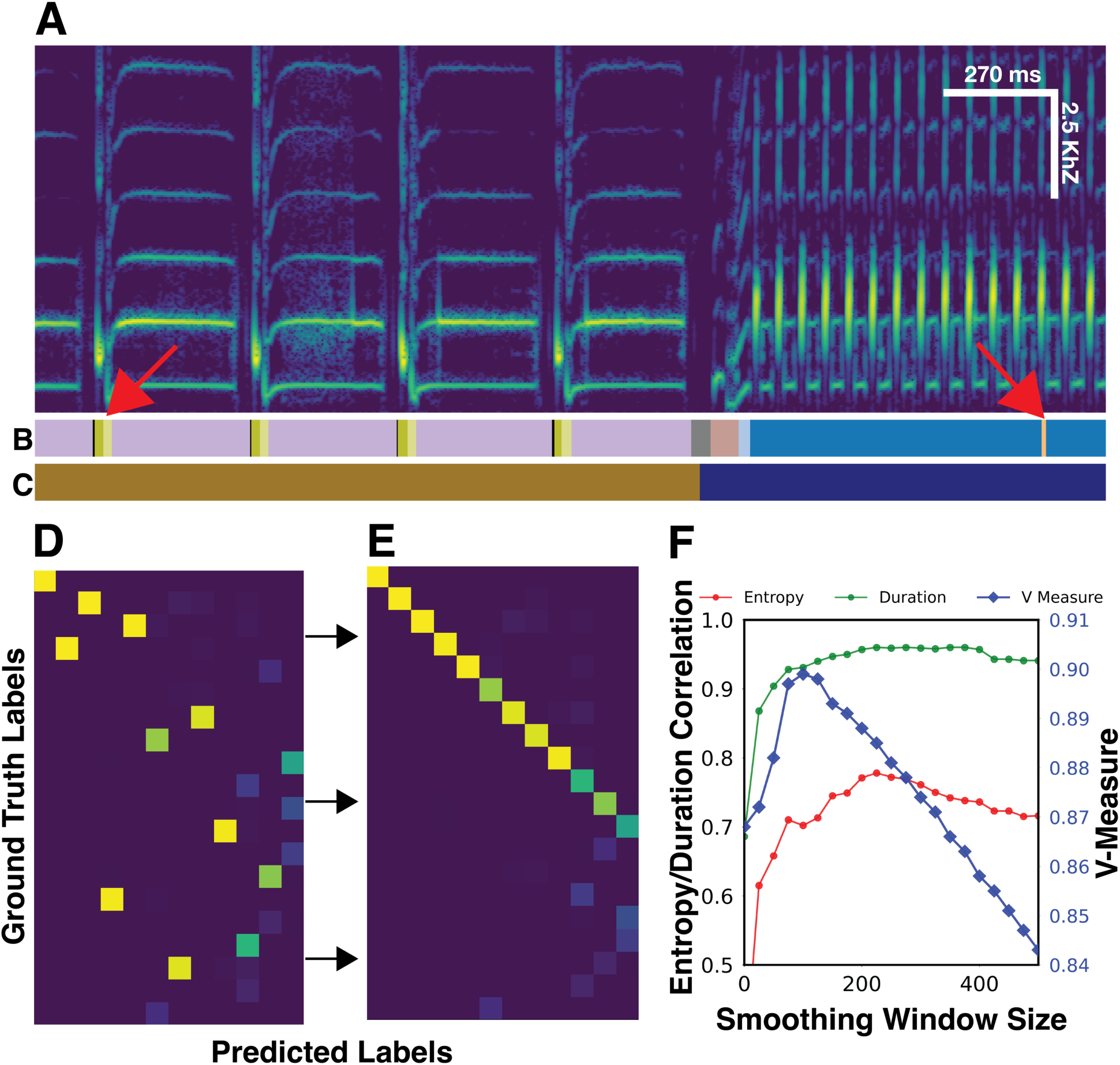
Comparing Human and Automated Labels for Sequence Analysis. **(A)** Spectrogram of canary song with corresponding labels: **(B)** HDBSCAN-generated clusters, and **(C)** ground truth human annotations. Red arrows indicate labeling discrepancies: syllable fragmentation (left) and spurious cluster insertions (right). **(D-E)** Confusion matrices comparing predicted and ground truth labels before **(D)** and after **(E)** alignment optimization. Misaligned or fragmented syllable predictions appear as off-diagonal elements, whereas unmatched machine-generated clusters occupy the lower rows of **(E)**. **(F)** Pearson correlations between HDBSCAN-derived and ground truth labels for entropy (▲), duration (●), and v-measure (▪) as a function of smoothing-window size. Optimal smoothing aligns human and model annotations at maximal correlation.

**Fig. 6.**
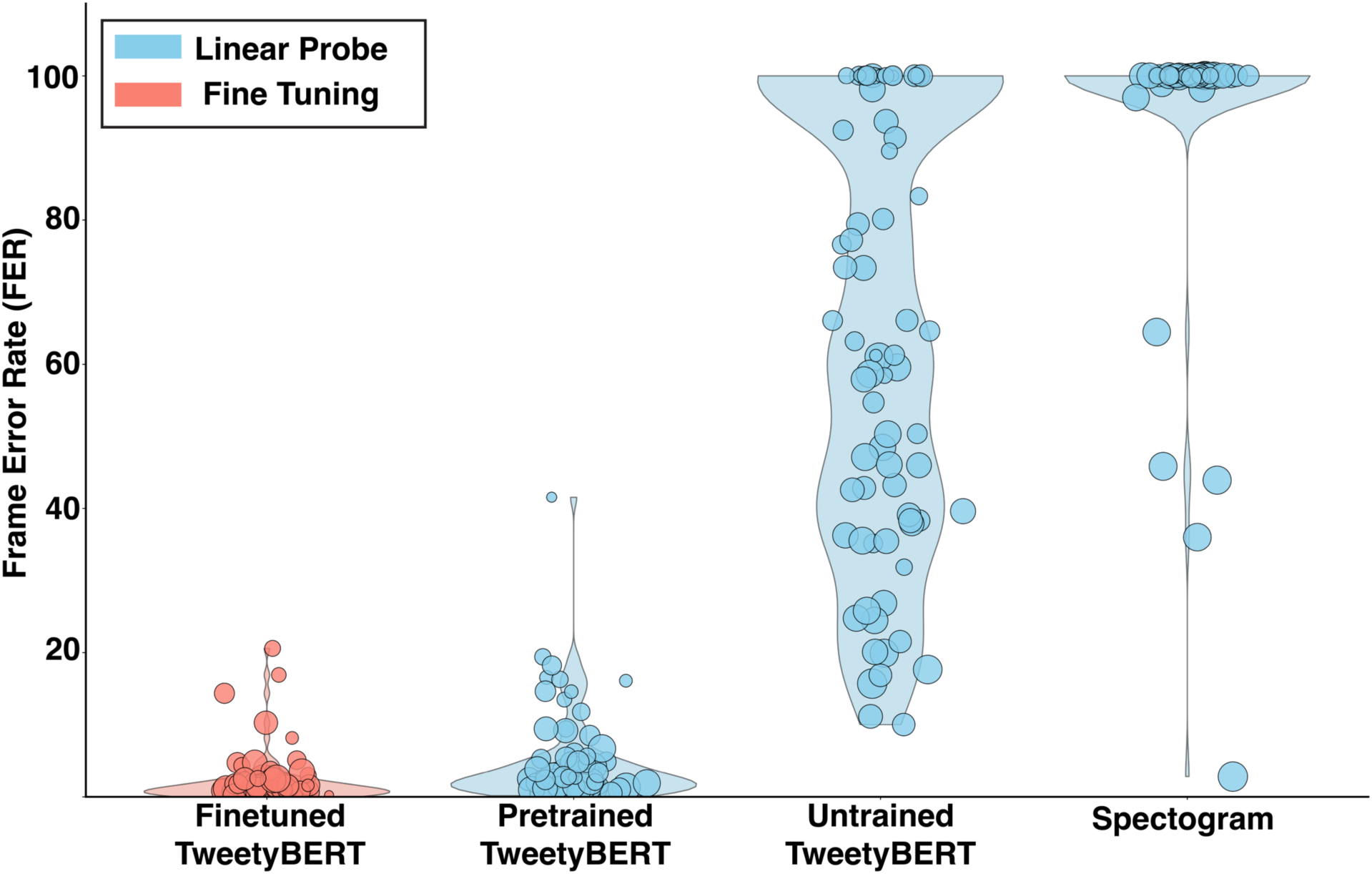
Evaluating TweetyBERT Embeddings Using Linear Probes. Frame error rates (FER) comparing finetuned TweetyBERT to linear probes on embeddings from pretrained TweetyBERT models, untrained TweetyBERT models, and spectrograms. Embeddings from pretrained TweetyBERT achieve near-finetuned accuracy, substantially surpassing those from untrained TweetyBERT or raw spectrogram features.

### Limitations and Potential Improvements

While many discrepancies are valid alternative conventions for song parsing, a small number clearly constitute errors. For example, the insertion of spurious cluster labels—often lasting just a few time bins—represents misclassification errors. Although these insertions are usually short in duration, they significantly impact downstream sequence analyses such as phrase durations, entropy measurements, and syntactic pattern analysis. These short errors can often be corrected through a post-processing step of temporally smoothing the syllable labels. This smoothing will be further discussed later.

### Mapping Machine to Human Labels

To further evaluate how well automated (machine) labels align with human annotations we identified the best matching machine labels for each human-labeled class. We constructed a co-occurrence matrix counting overlaps between predicted clusters and human labels, then used the Hungarian algorithm to derive an optimal one-to-one mapping that maximizes total overlap (Munkres, 1957). Since HDBSCAN typically discovers more clusters than exist in the ground truth labels, we grouped any unmatched machine labels into a separate noise or unmatched category.

### Introduction and Motivation for Smoothing

As mentioned earlier, automated cluster assignments occasionally contain spurious fluctuations or fragmented classifications, and these can significantly distort sequence measures such as phrase durations or transition entropy. To address this, we applied temporal smoothing—a procedure that assigns each time bin to the most frequently occurring state within a moving temporal window. This approach reduces brief misclassifications, stabilizes cluster assignments, and enhances the interpretability and reliability of sequence analysis metrics.

### Evaluation Metrics

With the machine to human label mapping or “Rosetta stone” in hand we characterized the match between human and machine annotations in three distinct ways. First to quantify classification accuracy, we introduced two variants of Frame Error Rate (FER). Matched-Only FER calculates error percentage considering only time bins assigned to matched clusters, thereby evaluating errors for recognized phrase types. Total FER counts all unmatched or misclassified time bins (including noise/unmatched categories), offering a comprehensive measure of overall classification accuracy. In addition to frame error rates, we compare higher order statistical measures for song between human and machine labels. Specifically, we focused on: phrase duration**—**the average length (in time bins) of each phrase type and phrase entropy**—**the variability of transition probabilities between states (Koparkar et al., 2024). We compute the correlation between phrase duration and phrase entropy computed by humans and machine, for each syllable in the data set. These measures can be quite fragile – even a single misclassified time bin can break a phrase into multiple pieces, so we sought to perform this measure with various degrees of data smoothing.

### Effect of Smoothing Window Size

We varied the temporal smoothing window from 0 to 500 time bins in increments of 25 time bins. Smaller windows (0–50 time bins) yielded more fragmented classifications, higher FER scores (matched only FER ≥ 6.76%, total FER ≥ 15.12%), and lower entropy correlations (r ≤ 0.658), despite achieving high V-measure scores (up to 0.882). Moderate smoothing windows (150–225 time bins) substantially improved overall performance, resulting in strong phrase-duration correlations (r ≥ 0.947), significantly improved entropy correlations (r ≥ 0.745), and lower error rates (matched only FER ≤ 4.99%, total FER ≤ 14.52%). Optimal performance was achieved at a 200 time bin window (∼540 ms), achieving the lowest matched-only FER (4.34%), a low total FER (13.97%), and the highest entropy correlation (r = 0.771), albeit with a slight decrease in V-measure (0.888 vs. 0.893 at 150 bins) (Supplementary Table 5 and Supplementary Fig. 6).

### Recommendations and Practical Implications

For canary song, we recommend a default smoothing window of approximately 200 time bins (∼540 ms) as this window size balances cluster fragmentation, classification accuracy, and entropy correlations effectively, and this duration corresponds roughly to the duration of most canary phrases (Markowitz et al., 2013). However, the ideal window size ultimately depends on analytical goals, such as minimizing frame error rates, maximizing entropy correlations, or preserving cluster accuracy as measured by V-measure. Practically, smoothing windows between 150 and 225 bins offer robust compromise solutions, effectively capturing essential syntactic structure while minimizing labeling errors.

### Motivation and Background

The previous analyses of TweetyBERT’s embeddings relied on dimensionality reduction generated by UMAP. While useful, UMAP is highly nonlinear, making it challenging to quantitatively assess learned representations. To complement the UMAP analysis, we turned to linear probes, a widely used method for evaluating self-supervised neural network embeddings without reliance on nonlinear dimensionality reduction techniques.

Linear probes are simple linear classifiers—a single matrix multiplication—applied directly to a network’s internal activations to predict class labels (Alain & Bengio, 2016). Linear probes cannot create new representations; they can only rotate or scale existing ones. Linear probes have demonstrated that self-supervised transformers encode sophisticated semantic and syntactic structures without explicit supervision (Coenen et al., 2019; Hewitt & Manning, 2019; Li et al., 2022). High performance of a linear probe indicates that the underlying latent spaces are meaningfully structured.

Here, we use linear probes to evaluate whether TweetyBERT’s self-supervised training produces emergent phrase-level representations of canary song. To apply the linear probe to the model, we simply train a linear classifier to reproduce human phrase labels, using the latent space vectors produced by TweetyBERT.

### Experimental Conditions

We examined 4 different models, trained to reproduce the human labels. For the TweetyBERT linear probe tests, we used outputs of the third attention sublayer as the basis for classification.

1. **TweetyBERT Fine Tuning:** first test is not a linear probe, but a full fine-tuning of all parameters of TweetyBERT to reproduce the human labels. This provided the best frame error rate that can be achieved with the TweetyBERT architecture if full supervised training is allowed.
2. **TweetyBERT Linear Probe:** probe was applied to the self-supervised (trained) TweetyBERT model. All parameters other than the linear projection were frozen after self-supervised pretraining.
3. **Untrained TweetyBERT Linear Probe:** probe was applied to the outputs of the untrained (randomly initialized) TweetyBERT model. All parameters other than the linear projection were frozen. This condition assesses inherent inductive biases of the untrained architecture.
4. **Spectrogram Linear Probe:** linear probe was applied directly to the raw spectrogram features, confirming that syllables are not linearly separable based on their single-time bin spectral features.

### Linear Probe Results

FER scores were derived from a held-out dataset not used during pretraining or linear probe training. On this unseen data, the pretrained TweetyBERT achieved a total FER of 2.5% (class-level standard deviation: 6.3%), approaching the fully supervised fine-tuned model’s (FER: 1.3%, class-level standard deviation: 3.5%), indicating that self-supervised learning alone produces near-optimal phrase-level representations. As expected, the untrained TweetyBERT performed worse (FER: 45.1%, class-level standard deviation: 30.0%), yet surpassed the raw spectrogram baseline (FER: 82.6%, class-level standard deviation: 15.8%). Since both the untrained model and the raw spectrogram classifier used identical 196-dimensional input spaces, the considerable performance advantage (∼38%) of the untrained transformer indicates that its architecture inherently generates latent variables that mix information from distinct time points, providing a boost in performance. This observation aligns with recent studies suggesting that transformers have intrinsic inductive biases beneficial for structured tasks (Zhong & Andreas, 2024), and in this sense, the untrained transformer functions analogously to a reservoir computer or liquid state machine (Jaeger, 2001; Maass et al., 2002).

### Implications for Architecture Selection

TweetyBERT’s randomly initialized architecture provides a meaningful boost in classification, and this suggests that linear probes of randomly initialized networks could be used to select hyperparameters or architectures with favorable inductive biases before large-scale training. The self-supervised pre-training further refines these initial representations, resulting in phrase-level embeddings that closely approximate supervised fine-tuned performance.

### Seasonal Vocal Plasticity in Canaries

Canaries are seasonal songbirds known to exhibit an annual cycle of song relearning. The underlying mechanisms of neural plasticity include large scale neural replacement that provides canary brains with newborn neurons capable of song relearning each fall (Nottebohm et al., 1986; Voigt & Leitner, 2008). To investigate how TweetyBERT captures these seasonal differences, we trained a new model combining spring and fall song data from two birds. From this model, we generated embeddings comprising approximately one million time bins per bird, divided evenly into four temporally distinct subsets (∼250,000 bins each): two subsets from the breeding (spring) and two from the non-breeding (fall) seasons. All spring songs were color-coded purple, and fall songs green. We note that the spring songs do not appear as well organized as prior UMAP representations of spring song generated by TweetyBERT. This is likely because the model combined spring and fall recordings which have different acoustic characteristics.

To further quantify embedding similarity, we constructed 300×300 binned heatmaps representing point densities in UMAP space, and quantified overlap of these densities using the Bhattacharyya coefficient. Within-season comparisons revealed high stability of these UMAP density plots for both canaries (canary 1: breeding = 0.938, non-breeding = 0.841; canary 2: breeding = 0.926, non-breeding = 0.930). However, between-season analyses showed moderate overlap for canary 1 (0.477) and lower overlap for canary 2 (0.119), highlighting more substantial seasonal vocal reorganization in canary 2.

These observations align with prior reports on seasonal vocal plasticity in canaries. The consistently high within-season similarity scores indicate stable song that has been termed “crystalized song” by researchers, whereas the lower between-season similarity indicates significant repertoire restructuring in the transition between breeding season and fall, a pattern visually evident in the UMAP embedding of canary 1’s vocalization. These observations in a small number of birds suggests that TweetyBERT embeddings may provide a useful latent space for a variety of song analysis tasks. Future studies will be needed to compare the performance of TweetyBERT relative to other unsupervised representations of song for qualitative analysis of latent spaces.

## Discussion

In this work, we introduced TweetyBERT, a self-supervised transformer model capable of automatically discovering the units of canary song without relying on human training data. Unlike many transformer models for human speech, TweetyBERT directly processes spectrograms without tokenizing the sound into discrete units. This results in a model with very few hyperparameters that need to be adjusted, and the approach should be applicable with minimal variation to a wide variety of species. In addition to clustering syllables with a good correspondence to human labels, the latent space generated by the model can be used to qualitatively “fingerprint” vocal repertoires as demonstrated with our analysis of seasonality in canary song. These latent space representations may prove to be useful in analysis of juvenile song, or other songs that are not as easily clustered as adult canary song. The initial findings reported here demonstrate the potential of self-supervised transformer architectures to significantly advance animal communication research.

### Limitations

Several practical challenges remain for broadly deploying self-supervised models like TweetyBERT in animal communication studies. Foremost an automated “song detector” is likely essential to isolate songs from cage noise or other background sounds prior to training. We have not examined whether TweetyBERT can be trained to parse song if cage noise or other environmental noises contaminate the recordings. If the model uses its representational capacity to predict these non-song acoustic elements, its performance on song will likely be reduced. In contexts where robust song detectors do not exist—particularly for niche or understudied species—developing custom song detectors introduces additional overhead.

Although TweetyBERT effectively matches human labels for most syllables, some discrepancies persist between automated clusters and human annotated labels. These differences in most cases reflect a difference of convention with human practice – such as splitting long multi-part syllables into distinct clusters or combining clusters that are not reliably separated. The field needs new methods to quantify the quality of automated clustering algorithms. If the machine learning provides a better parsing of song than humans do, how can we recognize this?

Some of the observed discrepancies in clustering suggest limitations of HDBSCAN’s spatial clustering method, and in particular its inability to distinguish partially overlapping acoustic trajectories. A history-dependent clustering approach, such as Autoregressive Hidden Markov Models (AR-HMMs), may improve cluster separation by incorporating sequential dependencies (Lin et al., 2024).

Systematic hyperparameter tuning can be a challenge in training large-scale audio models. Our current selection of hyperparameters (e.g., context length, masking strategies, or number and size of transformer layers) was guided by current best practices of parameter count to data ratios (Hoffmann et al., 2022), but we did not explore the model parameter space in any meaningful way. The use of linear probes applied to networks with random parameters may prove useful in guiding future model design. Still, current evaluation metrics depend on ground truth human annotations, which may not always be available or unbiased. Future unbiased measures of model performance could include decoder models that predict the spectral structure of song from the latent space, or from the discrete labels of the TweetyBERT model.

Computational limitations present another barrier. UMAP dimensionality reduction becomes computationally prohibitive when processing large datasets comprising millions of data points, restricting scalability and reproducibility. While these limitations do not invalidate insights derived from smaller-scale analyses, they hinder the broader deployment of TweetyBERT for large-scale data analysis. Approaches to mitigate this limitation of UMAP include the development of decoders trained to predict the cluster labels generated by the TweetyBERT model, so that the UMAP step can be avoided in large-scale inference tasks.

### Future Directions

We chose an input data representation consisting of high-resolution spectrograms, computed using parameters commonly applied in the field – similar spectrogram timescales have historically been used for analysis of canary song. It will be valuable to compare the performance of TweetyBERT with spectrograms of distinct timescale and bandwidth. Is it necessary to know in advance which spectrogram parameters work well for a given species, or can the TweetyBERT architecture perform equally well over a range of parameter settings? We anticipate that choice of spectrogram parameters will not be critical, so long as the temporal resolution of the spectrogram can capture the shortest syllables in the song.

Models such as wav2vec2.0, Whisperseg, and HuBERT/AVES process raw waveforms but typically impose fixed-duration convolutional kernels in the front end (typically 20ms for human speech) (Baevski et al., 2020; Gu et al., 2023; Hagiwara, 2022; Hsu et al., 2021). TweetyBERT’s fundamental timescale is fixed at 2.7ms. Recent advances in NLP, such as the Byte Latent Transformer (BLT), dynamically allocate smaller tokens in information-rich regions, bypassing fixed tokenization entirely (Pagnoni et al., 2024). Adapting these strategies to bioacoustics data may facilitate modeling diverse vocal repertoires across multiple species.

Pixelwise reconstruction of missing spectrogram patches is fragile in that small variations in the spectral or temporal structure of predicted spectrogram fragments can lead to significant variations in the reconstruction score. Similar limitations held back the effective use of transformer models in image and video tasks which motivated the recent development of latent-space predictive architectures such as Joint-Embedding Predictive Architectures (JEPA) (Assran et al., 2023; Fei et al., 2023) could improve the objective function that guides self-supervised learning. These more advanced models may be necessary to achieve robust performance with more complex vocalizations such as juvenile song, or more complex acoustic environments such as environmental recordings in the wild.

Future work may explore models incorporating internal dimensionality reduction, eliminating reliance on external methods like UMAP, as demonstrated successfully in neural activity embeddings (Schneider et al., 2023). Furthermore, extending transformer context windows beyond the 2.7s used here could improve performance by capturing longer-range dependencies in vocal sequences (Markowitz et al., 2013).

### Broader Implications of Machine Learning in Animal Vocalization Data

Our findings highlight the potential for self-supervised models like TweetyBERT to enhance research in animal communication. While we have focused on models applied to single individuals singing alone, a range of groups are working on methods to capture social conversations (Rüttimann et al., 2023; Schulthess et al., 2023). Models like TweetyBERT can be trained on multi-individual conversations, and masked audio can be predicted from the surrounding context of the entire conversation, rather than acoustic context of single individuals. Latent space representations of vocal communication in social contexts could reveal how the structure of the vocalizations depend on the identity of the conversational partner, or what that partner said. Analysis of social conversations in songbirds and other vocal communicators like parrots or dolphins has already begun to reveal previously hidden meanings in vocal behavior (Balsby et al., 2012; King & Janik, 2013), with recent unsupervised clustering and latent-space modeling of sperm whale communication providing clear examples of this emerging science (Andreas et al., 2022; Begus et al., 2023; Sharma, Gero, Payne, et al., 2024; Sharma, Gero, Rus, et al., 2024).

Thus far, TweetyBERT has been applied to high quality audio recordings gathered in a laboratory setting. Field recordings include higher levels of background noise and more complex echoes, suggesting future challenges to applying this method to environmental audio. Still, successful deployment on field recordings could enable transformative ecological applications. Automated song analyses would significantly enhance long-term monitoring of wild songbird populations, revealing how environmental factors shape vocal behavior or dialect formation. Furthermore, automated song identification could become a critical conservation tool, aiding in the assessment of ecosystem health and tracking individual animals within populations. Ultimately, widespread adoption of these technologies will deepen human understanding and appreciation of animal communication and behavior. Across a wide range of animal species from songbirds to dolphins, the contextual information so easily captured by the transformer architecture, may provide the essential advance needed to accelerate the scientific study of animal communication.

## Materials and Methods

### Animal Subjects and Housing

This study utilized two datasets of adult male American Singer canaries (*Serinus canaria*): the TweetyNET dataset and a newly collected seasonal dataset. The TweetyNET dataset contained recordings and syllable-level annotations from three birds and was previously described in detail (Cohen et al., 2022). The seasonal dataset comprised recordings from two birds across both breeding and non-breeding seasons.

All procedures followed the protocol approved by the University of Oregon Institutional Animal Care and Use Committee (IACUC). For the seasonal dataset, canaries were sourced from Maryland Exotic Birds (Pasadena, MD) and housed at the University of Oregon’s Terrestrial Animal Care Services (TeACS). Birds were maintained in flight cages on a light cycle synchronized to seasonal changes. Bird seed (Kaytee Supreme Seed), pellets (Lafeber’s Premium Canary pellets), high-calcium grit, and fresh water were offered ad libitum. For experimental recordings, birds were singly housed in cages (Model C-1510, Kings Cages International, Ft. Lauderdale, FL) placed inside environmental chambers (Omnitech Electronics Inc., Columbus, OH). Sound attenuating foam (Sonex Classic, Pinta-Acoustic, Minneapolis, MN) was used to line the interior of the chambers in an effort to minimize external noise. Birds had *ad libitum* access to Kaytee Supreme Seed and Lafeber’s Premium Canary Pellets, and freshwater. Enrichment for isolated birds included socialization (1-2hr/day, 3-5x/week) in addition to treats (dried mealworms, dried fruit, crinkle paper, and mixed greens).

### Audio Recording and Data Acquisition

The TweetyNET dataset includes recordings from three canaries, collected under conditions detailed in the TweetyNET paper (Cohen et al., 2022).

For the seasonal dataset, recordings were made using an omnidirectional microphone (Audio-Technica AT803) positioned above each cage. The audio was routed through an M-Audio 8 pre-amplifier and captured using Sound Analysis Pro 11 software. Recordings were conducted at 44.1 kHz (single channel), generating WAV files.

### Spectrogram Generation and Preprocessing

Spectrogram generation parameters were adapted from the TweetyNET paper’s established methodology. Raw audio recordings were processed using a 5th-order elliptic high-pass filter (0.2 dB ripple, 40 dB stopband attenuation, 500 Hz cutoff) to remove low-frequency noise that could interfere with song analysis. Short-Time Fourier Transform was then applied using a Hann window with a 1024-point FFT and 119-sample hop length, followed by amplitude-to-dB conversion referenced to maximum amplitude. These parameters were chosen to optimize the time-frequency resolution trade-off for capturing fine temporal structure of syllables while maintaining sufficient frequency resolution for spectral analysis. Spectrogram generation was restricted to time intervals marked as containing songs by the song detector (Supplementary Fig. 2). Spectrogram processing was performed using librosa (McFee et al., 2015) and soundfile (https://github.com/bastibe/python-soundfile).

### Model Architecture (TweetyBERT)

TweetyBERT is a self-supervised, transformer model designed to learn representations of canary song directly from spectrogram inputs. The model processes spectrograms spanning 196 frequency bins by 1000 time bins, corresponding to 2.7 seconds of audio. Although a 2.7-second context window does not encompass entire canary songs, it sufficiently captures between one and three complete song phrases. We avoided using longer segments, such as entire songs, due to the quadratic memory requirements inherent in transformer architectures. This segment length also aligns historically with prior supervised deep learning methods, where optimal segments were approximately 1 second long (Cohen et al., 2022). Input spectrogram segments were trimmed from the original 512 frequency bins down to 196 bins, preserving the vocal frequency range of canaries while reducing memory usage.

The overall TweetyBERT architecture sequentially integrates the following components: (1) spectrogram masking, (2) a convolutional neural network (CNN) front-end for local feature extraction, (3) linear projection of CNN outputs into a Transformer-compatible embedding space, (4) stacked Transformer encoder blocks for modeling temporal dependencies and feature encoding, (5) linear projection back into spectrogram space, and (6) computation of a masked reconstruction loss (mean squared error).

The convolutional front-end contains four convolutional layers (Conv1 with 32 channels, Conv2–Conv4 with 64 channels each), each followed by GELU activation and max-pooling operations. Specifically, convolutional layers utilize a kernel size of 5×5, padding of 2, and are followed by max-pooling layers with kernel sizes of (2,1) to reduce dimensionality selectively along the frequency axis while preserving temporal resolution.

The Transformer encoder is the central component of TweetyBERT, consisting of four identical encoder blocks adapted from the original BERT architecture, incorporating enhancements to improve performance and training stability. Each encoder block comprises a multi-head self-attention layer with four attention heads (head dimension = 49, total embedding dimension = 196), followed by a feed-forward neural network (FFN) with a hidden layer dimension of 768. The Transformer blocks incorporate relative learned positional encodings, enabling the model to represent contextual relationships between time bins based on their relative positions rather than absolute locations, thus providing translation invariance across the sequence (Shaw et al., 2018). We further adapted the original BERT design by applying pre-layer normalization to improve gradient stability and training convergence, and included GELU activation functions within the FFN layers to provide smooth nonlinear transformations (Hendrycks & Gimpel, 2016; Xiong et al., 2020). Qualitatively, we found these modifications substantially improved training speed, convergence behavior, and overall stability. The complete TweetyBERT model (CNN front-end plus Transformer encoder) contains approximately 2.5 million parameters.

### Training Protocol and Preprocessing

For pre-training TweetyBERT, the dataset was partitioned into an 80/20 train-test split at the song level, ensuring that entire songs were exclusively assigned to either the training or test set. This strategy prevented data leakage by ensuring the model never encountered fragments of the same song in both sets. Two distinct models were trained: one on the TweetyNET dataset, comprising ∼15 hours of song for training and ∼4 hours in a holdout set used for early stopping and downstream analysis, and another for seasonality analysis, trained with ∼20 hours of song and a holdout set of ∼5 hours. The TweetyNET dataset model was trained for 26,500 optimization steps, while the seasonality model underwent 35,500 optimization steps, with each optimization step corresponding to one batch.

Raw spectrogram inputs underwent several preprocessing steps. The frequency dimension was truncated to the range [20, 216] frequency bins [860hz, 9288hz], retaining the frequencies most relevant to canary vocalizations. Each spectrogram was z-score normalized across the entire frequency-time matrix (not column wise z-scoring, but image-wise), stabilizing input distributions and improving training stability. During training, the data loader dynamically extracted random 1000 time bin windows (∼2.7 seconds) from larger spectrograms. This prevented the network from memorizing fixed segments and ensured diverse input exposure across batches. To enhance generalization, random frequency shifts of ± 50 (∼2kHz) frequency bins were applied to the spectrogram image during training, with shift values sampled uniformly from 0-50 frequency bins. This introduced significant variation in frequency content that was beyond the natural frequency variability of adult canary song syllables. For spectrograms shorter than 1000 time bins, zero-padding was applied to maintain consistent input dimensions.

A masked prediction objective guided the model’s self-supervised learning. Twenty-five percent of the input spectrogram was randomly occluded by contiguous segments of varying lengths, sampled uniformly from 0 to 250 time bins (0 – ∼675 ms). Masked regions were replaced with zeros, encouraging the model to infer missing values based on the surrounding context. To increase the per training step training speed, identical masks were applied across all batch elements per optimization step.

### Loss, Optimization, and Early Stopping

The training objective minimized a mean squared error (MSE) loss, computed exclusively over the masked regions. Optimization was performed using the Adam optimizer with a learning rate of 3e-4 and a batch size of 42 (Kingma & Ba, 2014). The training loop utilized mixed-precision training with torch.cuda.amp and GradScaler to reduce memory consumption and increase training speed (Micikevicius et al., 2018). Validation was conducted at regular intervals (every 500 steps), with the validation loss smoothed using a 1000-step moving average to mitigate noise in performance evaluations.

An early stopping criterion was employed to ensure efficient convergence. Training terminated if the smoothed validation loss did not improve for eight consecutive validation checks. This mechanism prevented excessive overfitting and halted training once the model had converged.

### Embedding Generation and Dimensionality Reduction

To generate TweetyBERT embeddings, up to 1 million spectrogram time bins (∼45 min of song) were processed (Fig. 3 & 7). Songs from each bird were segmented into 1000 time bin windows matching the model’s context window size, with zero-padding applied to the final segment as necessary. Segments were batched and passed through the model, collecting attention activations from the third transformer layer (Supplementary Table 7), chosen due to its high V-measure scores. The resulting representations were concatenated to reconstruct song-level features, ultimately yielding 1 million time bins of neural activations. These 196-dimensional activations were then reduced to 2D via UMAP (cosine distance, 200 neighbors, minimum distance=0.1, seed=42). For spectrogram embeddings, UMAP was applied directly to z-scored spectrogram columns.

**Fig. 7.**
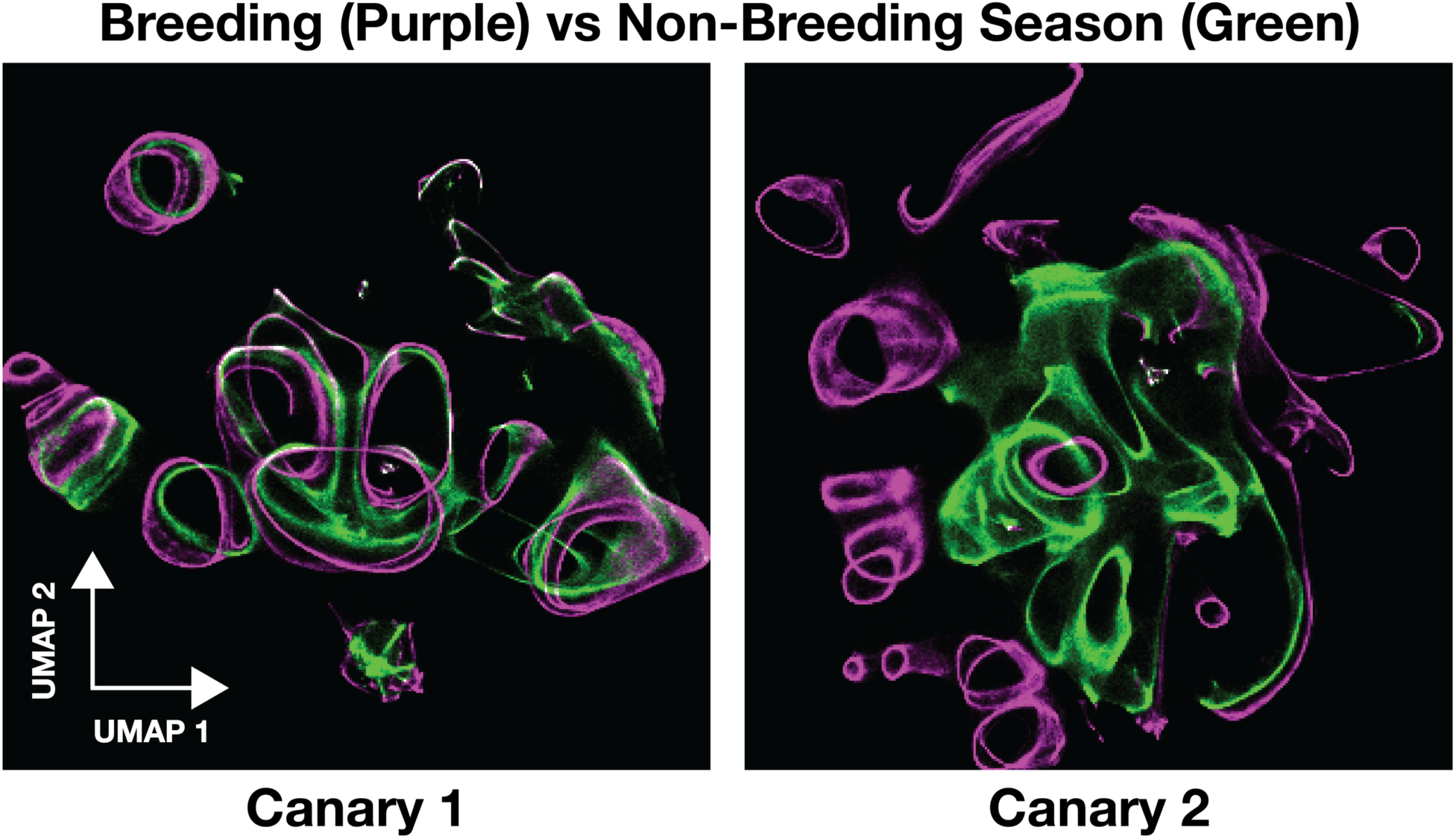
TweetyBERT embeddings vary by season of the year. UMAP embeddings of breeding season song (purple) and fall season song (green) of two canaries. Regions of overlap appear in white.

For temporal sequence analysis, clustering optimization, and V-measure calculations, only unseen songs from the holdout set were used for embedding generation, providing a measure of the model’s generalization performance. We additionally used data “folds” because embedding the entire test set at once would exceed manageable data limits for a single UMAP. Breaking the data into multiple folds (ranging from 288,876 to 685,969 time bins, averaging ∼442,502) also allowed us to construct statistical measures (e.g., V-measure) across random dataset breakdowns for each bird, thus capturing the variability in performance within and between folds.

### Syllable-to-Phrase Label Conversion

When necessary, we converted the existing syllable-level ground truth labels from the TweetyNET dataset into phrase-level labels by replacing inter-syllabic silent bins with their nearest adjacent syllable label. In case a of a tie, we took the syllable label earlier in the song. For silent regions at the beginning or end of a file (i.e., where no bounding label was present on one side), the silence bins were merged with the available adjacent syllable.

### Clustering with HDBSCAN

TweetyBERT embeddings were clustered using the HDBSCAN implementation provided by (McInnes et al., 2017), with minimum samples=1 and a minimum cluster size of 5000.

### Majority Vote Smoothing

After clustering with HDBSCAN, we applied a majority-vote smoothing algorithm to reduce transient fluctuations in cluster assignments. Specifically, for each labeled sequence of length *N*, we took a sliding window of length *w* around each position *i* and replaced the label at position *i* with the most frequent label in that window. In the event of a tie, the algorithm selected the label that appeared first in the window. At the start and end of sequences, window sizes were dynamically adjusted to fit the available data.

### Replacing Noise with Nearest Neighbor

HDBSCAN designates outlier points as noise. In practice, only a small fraction of time bins were labeled as noise by HDBSCAN. To preserve potentially meaningful data, we reassigned these noise-labeled points to the nearest valid cluster within the same sequence of labels. Specifically, for each noise-labeled bin, we searched leftward and rightward to locate the nearest labeled bins, measured the distance in time bins, and adopted the label of the nearest neighbor. If both neighbors were equidistant, we defaulted to the left label. In rare cases where neither side contained a valid label, the original noise label remained unchanged. This procedure was applied to the V-measure, correlation, and smoothing analyses.

### V-measure

V-measure is a widely used metric for evaluating clustering performance, calculated as the harmonic mean of two complementary components: homogeneity, which assesses how well clusters contain members of a single ground truth class, and completeness, which evaluates how well each ground truth class maps to individual clusters (Rosenberg & Hirschberg, 2007) (Pedregosa et al., 2012). The V-measure ranges from 0 to 1, where 1 indicates perfect clustering and 0 indicates complete disagreement between clusters and ground truth labels.

### Temporal Label Mapping and Refinement

Temporal label mapping and refinement: A shared-area matrix, *M*, was constructed such that each entry *M*₍ᵢ,ⱼ₎ represented the number of frames in which ground truth label *i* co occurred with predicted label *j*. A noise label was explicitly included in both ground truth and predicted sets to capture unmatched frames. *M* was then column normalized. Subsequently, the Hungarian algorithm (Munkres, 1957) was applied to the normalized matrix, yielding an optimal one-to-one assignment pattern between ground truth labels and predicted clusters that maximized diagonal alignment. Importantly, this approach matches each ground truth label exclusively to the single best predicted cluster, inherently leaving lower-ranked or secondary candidate clusters unmatched. Predicted clusters not mapped by this assignment were reassigned to the noise label, designating them as noise or unmatched. Frame-error rates (FER) were computed in two ways: a matched FER was calculated by comparing predicted labels with mapped ground truth labels for frames with valid mappings (excluding those labeled as noise), and an unmatched FER was derived by treating all time bins corresponding to unmapped predicted clusters (noise) as errors. Phrase-level correlation analyses (entropy and length metrics) were computed only for matched label pairs, as these analyses inherently required mapped pairs between ground truth and predicted clusters.

### Phrase Entropy and Duration

For each mapped pair of ground truth and HDBSCAN-predicted phrase labels, we computed the following metrics for both human (ground truth) and automated (predicted) annotations:

### Phrase Transition Entropy

For each phrase, we recorded how often it transitioned to each other phrase in the dataset. We then used Shannon entropy to measure the unpredictability of these transitions, averaging across all occurrences of the phrase. Higher entropy implies a more varied set of possible “next” phrases, whereas lower entropy indicates more predictable transitions (Koparkar et al., 2024).

### Phrase Duration

Phrase duration was defined as the average length of uninterrupted time bins assigned to each phrase label. Specifically, for each labeled phrase, we identified continuous runs of that phrase in the data and computed their mean duration.

### Weighted Pearson Correlation

To compare how well the automated (predicted) measures align with the human (ground truth) measures at the phrase level, we computed a weighted Pearson correlation. Each phrase received a weight proportional to its frequency of occurrence, ensuring that commonly used phrases had a greater influence on the correlation value. This approach quantifies agreement between ground truth and predicted metrics—such as transition entropy or duration—while giving more emphasis to phrases that appear more often.

### Linear Probe Analysis

The linear probe consisted of a single linear layer mapping 196-dimensional embeddings to phrase probabilities. For the pre-trained and untrained conditions, the probe was applied to the third attention layer, while for the fine-tuned condition, it was applied to TweetyBERT’s final linear deprojection layer. For the spectrogram condition, the probe was applied directly to raw spectrogram features. The linear probe was trained on the training set using Adam optimization (Kingma & Ba, 2014), with learning rates of 1e-2 (linear probe, spectrogram, untrained) and 3e-4 (fine-tuned); these values were crucial for convergence. Mixed-precision training (Micikevicius et al., 2018) and a batch size of 42 were used to reduce memory consumption and accelerate training. Frequency shifting augmentation or masking was not applied during training or evaluation of the linear probe. Early stopping with patience of six evaluation intervals (25 batches per interval) and a moving average of validation loss over 1000 steps were implemented to stabilize training and prevent overfitting. Training was capped at 5000 batches.

Accuracy was measured as frame error rate (FER), calculated as the percentage of mismatched time bins between linear probe predictions and ground truth labels. FER was computed as the average across the three labeled birds from the TweetyNET holdout dataset. The pretrained and fine-tuned models shared the same TweetyBERT backbone, trained on the TweetyNET dataset, while the untrained model was initialized randomly (He et al., 2015).

### Seasonal Embedding Analysis

TweetyBERT embeddings were used to characterize seasonal differences in canary song structure across breeding and non-breeding seasons. For each bird, song data from approximately one million time bins were projected into a common two-dimensional latent embedding space (UMAP coordinates). The resulting embeddings were divided equally into four distinct temporal groups: two groups from the breeding season and two groups from the non-breeding season.

Specifically, for canary 1, the recording dates for each group were:

- **Breeding Season 1:** May 28, 2024 – May 29, 2024
- **Breeding Season 2:** May 30, 2024 – May 31, 2024
- **Non-Breeding Season 1:** September 13, 2024 – September 21, 2024
- **Non-Breeding Season 2:** September 22, 2024 – September 27, 2024

For canary 2, the corresponding dates were:

- **Breeding Season 1:** May 30, 2024 – June 1, 2024
- **Breeding Season 2:** June 2, 2024 – June 3, 2024
- **Non-Breeding Season 1:** September 13, 2024 – September 16, 2024
- **Non-Breeding Season 2:** September 17, 2024 – September 20, 2024

The embeddings generated for each group were then aggregated into 2D histograms (heatmaps) using a 300×300 binning scheme. This process yields discrete probability distributions over the embedding space. Seasonal changes were quantified using the Bhattacharyya coefficient, a measure of overlap between probability distributions.

To assess song stability and seasonal changes, we compared:

- **Within-season stability:** The average Bhattacharyya coefficient between same-season groups (e.g., Breeding Season 1 vs. Breeding Season 2, Non Breeding Season 1 vs. Non Breeding Season 2).
- **Between-season differentiation:** The average Bhattacharyya coefficient between groups drawn from different seasons (e.g., Breeding Season 2 vs. Non Breeding Season 1).

### Software Packages

All neural network models, including TweetyBERT and the linear probes, were implemented and trained using PyTorch (Paszke et al., 2019). Spectrogram generation utilized Librosa (McFee et al., 2015) and Soundfile libraries. Clustering was performed using the HDBSCAN implementation of (McInnes et al., 2017), dimensionality reduction was achieved using UMAP (McInnes et al., 2018), and clustering performance was evaluated using V-measure from scikit-learn (Pedregosa et al., 2012). Numerical operations and array manipulations were performed using NumPy (Harris et al., 2020). Figures and visualizations were created using Matplotlib (Hunter, 2007) and Seaborn (Waskom, 2021).

### Computational Resources

All model training and analyses were performed using PyTorch on a system equipped with an NVIDIA GeForce RTX 4090 GPU (24GB VRAM), CUDA 12.3, driver version 545.23.08, and an AMD Ryzen threadripper pro 5955wx processor running Ubuntu 22.04.4 LTS with 128GB of RAM. Approximately 2TB of storage supported the extensive intermediate computations required. Parallel CPU computation accelerated spectrogram generation and feature extraction.

### Code Availability

TweetyBERT is available at https://github.com/georgevenven/tweety_bert. The TweetyNET song detector is available at https://github.com/georgevenven/tweety_net_song_detector.

### Data availability

The TweetyNET dataset used for model training is publicly available via Dryad (https://datadryad.org/stash/dataset/doi:10.5061/dryad.xgxd254f4). The seasonal dataset, trained model weights, and intermediate analysis files are available from the authors upon request.

## Acknowledgements

We would like to thank Diana Ostojich and Ananya Kapoor for providing edits and feedback. This work was funded by NIH R01NS118424.

---

**Supplementary Fig. 1.**
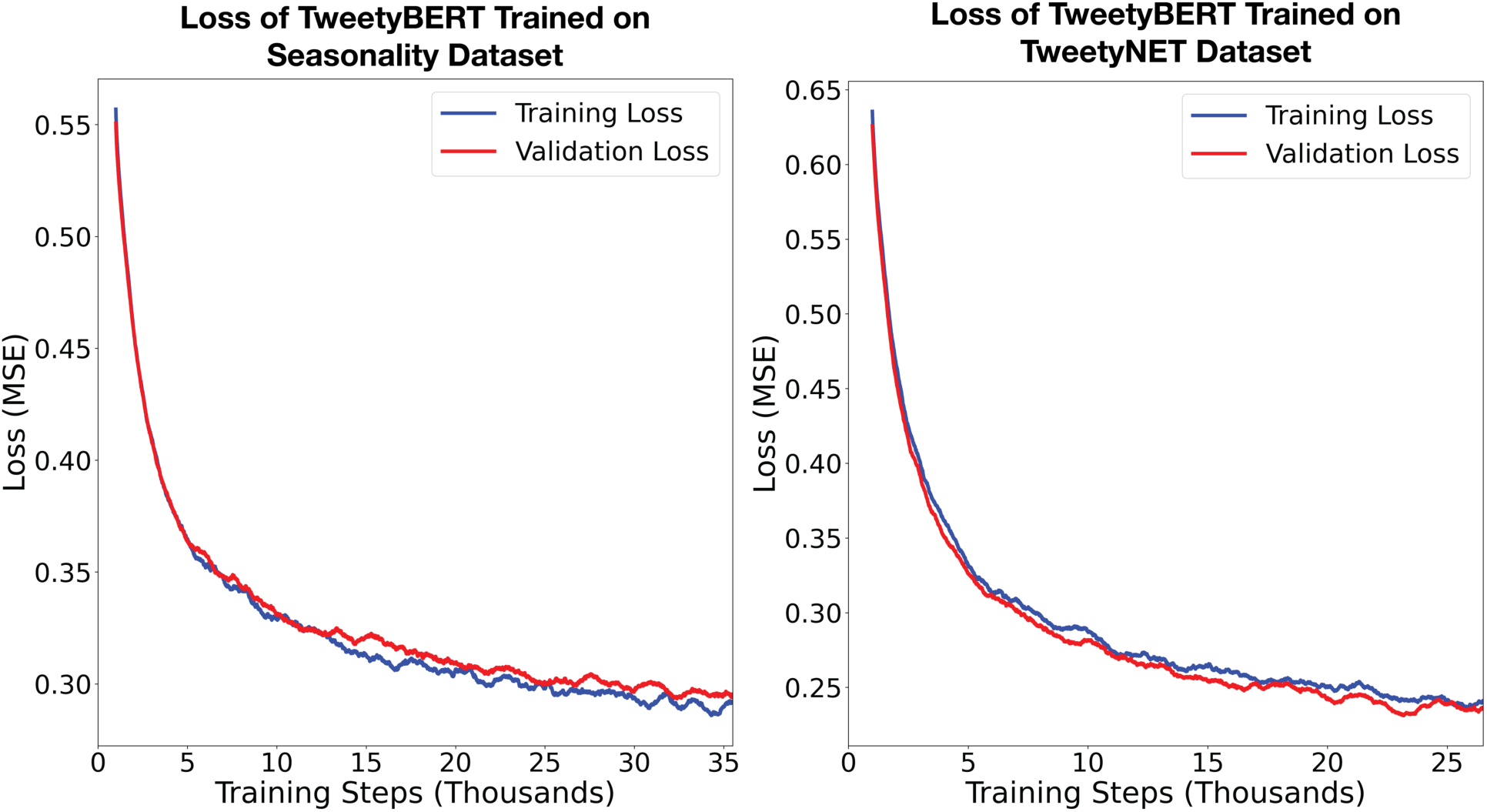
TweetyBERT MSE Loss vs Training Steps. Training and validation loss curves (mean squared error, MSE) for TweetyBERT trained separately on the Seasonality and TweetyNET datasets. Both training (blue) and validation (red) losses exhibit stable convergence, indicating effective self-supervised learning of masked spectrogram reconstructions. The higher final MSE observed for the Seasonality dataset likely reflects greater acoustic variability between breeding and non-breeding vocalizations, consistent with biologically driven seasonal vocal plasticity.

**Supplementary Fig. 2.**
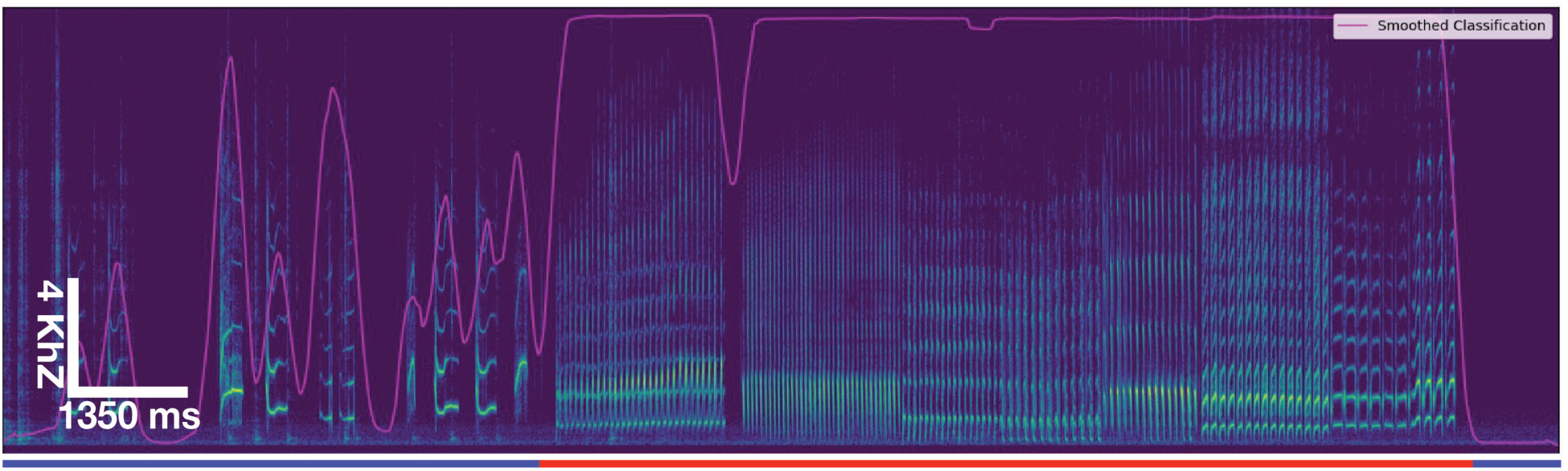
Song Detector. Spectrogram of a canary recording containing both song and non-song segments. The purple line represents the post-processed smoothed probabilities of each time bin being classified as canary song. The bottom bar displays the final Boolean classification of song (red) vs. non-song (blue) after post-processing. Notably, transient spikes around 1000 time bins—which are canary calls rather than song—are excluded from being labeled as song. The segmentation effectively distinguishes structured song from background noise, calls, and silent intervals, ensuring that only sustained vocal sequences are labeled as song. This figure represents a typical example of the classifier’s performance across the dataset.

The song detector used in this project is a lightweight variant of the TweetyNET architecture, designed to efficiently identify canary songs in spectrogram data. This model combines a convolutional front end with a bidirectional LSTM stack, maintaining a small hidden state size of 32 to optimize for speed and scalability. The song detector outputs a probability for each time bin, indicating the likelihood that the bin contains canary song. Ground truth labels are created by an expert human marking the start and stop of song. Spectrograms were generated using preprocessing methods previously described. During training, the model achieved a frame error rate (FER) of 2–3% on a test set containing 47 minutes of non-song and 27 minutes of song, demonstrating reliable detection of vocalizations versus background noise. The training was conducted with a learning rate of 3e-4, and the model was trained on 169 minutes of non-song and 107 minutes of song collected across a diverse set of recording conditions.

Post-processing plays a crucial role in refining the detector’s output and ensuring reliable song annotations. While the raw FER is low, direct predictions can suffer from over segmentation, splitting continuous songs into fragments, or under segmentation, incorporating non-song noise into detected segments. The post-processing pipeline mitigates these issues by smoothing predictions, applying thresholds to eliminate short and noisy segments, and padding detected boundaries to ensure no valid song content is truncated. Although padding can introduce brief silences at segment edges, this tradeoff is acceptable as this has negligible performance impacts on the training of TweetyBERT.

It is important to note that the model’s performance depends on the similarity between the training and deployment environments. Since the training data was manually annotated and collected in a consistent seasonal period and recording setup, the detector performs well within this context. However, deployment in different conditions may require retraining or fine-tuning to maintain accuracy.

**Supplementary Fig. 3.**
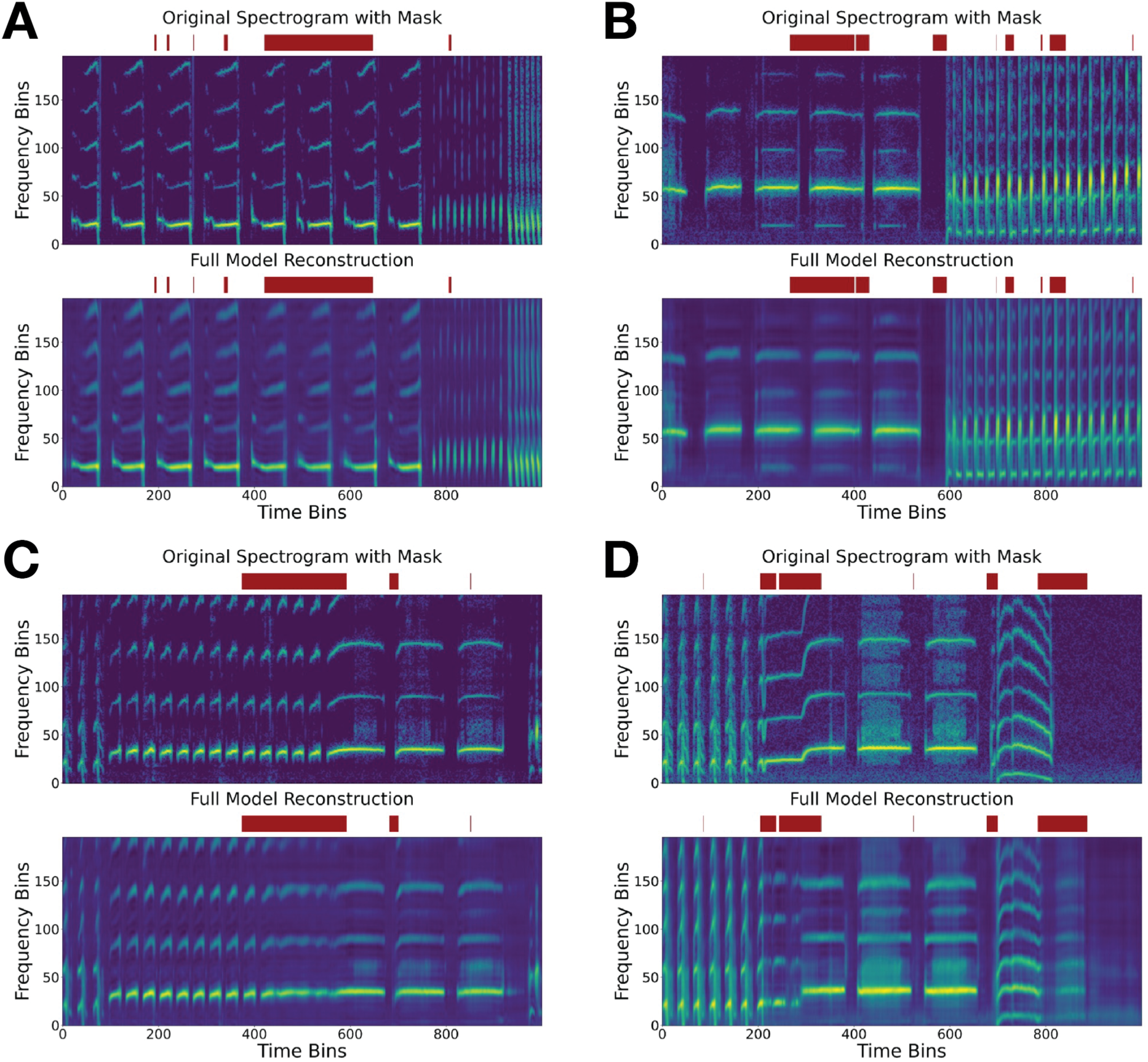
Montage of masked spectrogram predictions from the TweetyBERT training phase. Each panel consists of a masked spectrogram fed to TweetyBERT (top) and the model’s predicted reconstruction (bottom). Red bars indicate masked regions that were not visible to the model during this prediction. **(A, B)** Depict accurate reconstructions of masked segments, demonstrating the model’s capability to infer detailed acoustic structures from context. **(C, D)** Illustrate mild reconstruction errors where the model produces hybrid syllable morphologies by combining two syllable classes. These errors arise when critical contextual cues are obscured by masking, causing ambiguity in syllable prediction.

**Supplementary Fig. 4.**
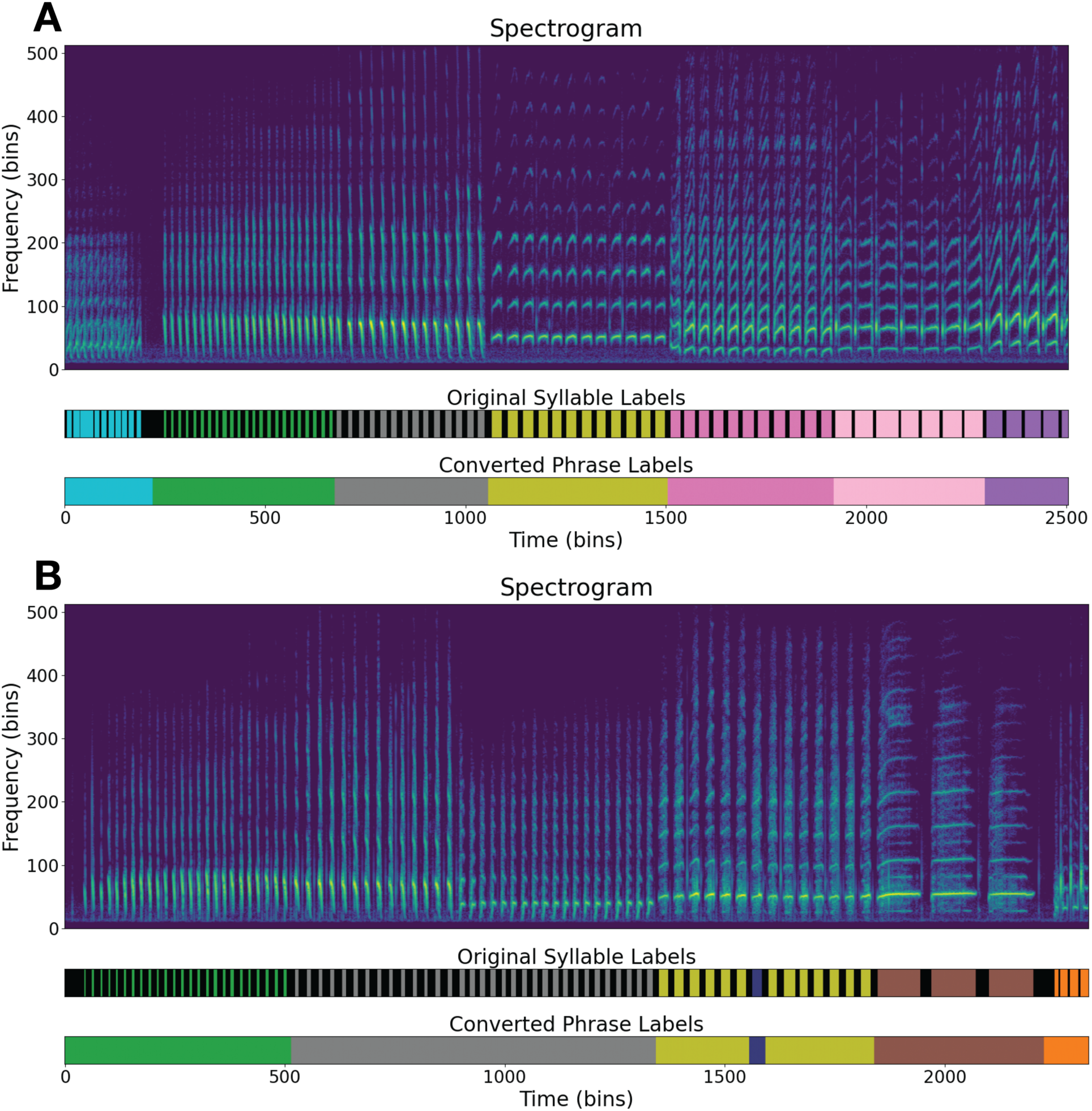
Examples of conversion from syllable-level to phrase-level labels. **(A, B)** Spectrograms of complete canary songs, each shown with original ground truth syllable-level annotations (top, silences in black) and their corresponding phrase-level labels after conversion (bottom). During conversion, silent intervals between syllables were reassigned to their nearest neighboring syllable labels. Panel **(B)** highlights two annotation errors in the original ground truth labels: a mislabeled syllable insertion (purple within yellow) and two distinct phrases incorrectly labeled as identical (gray).

**Supplementary Table 5.**
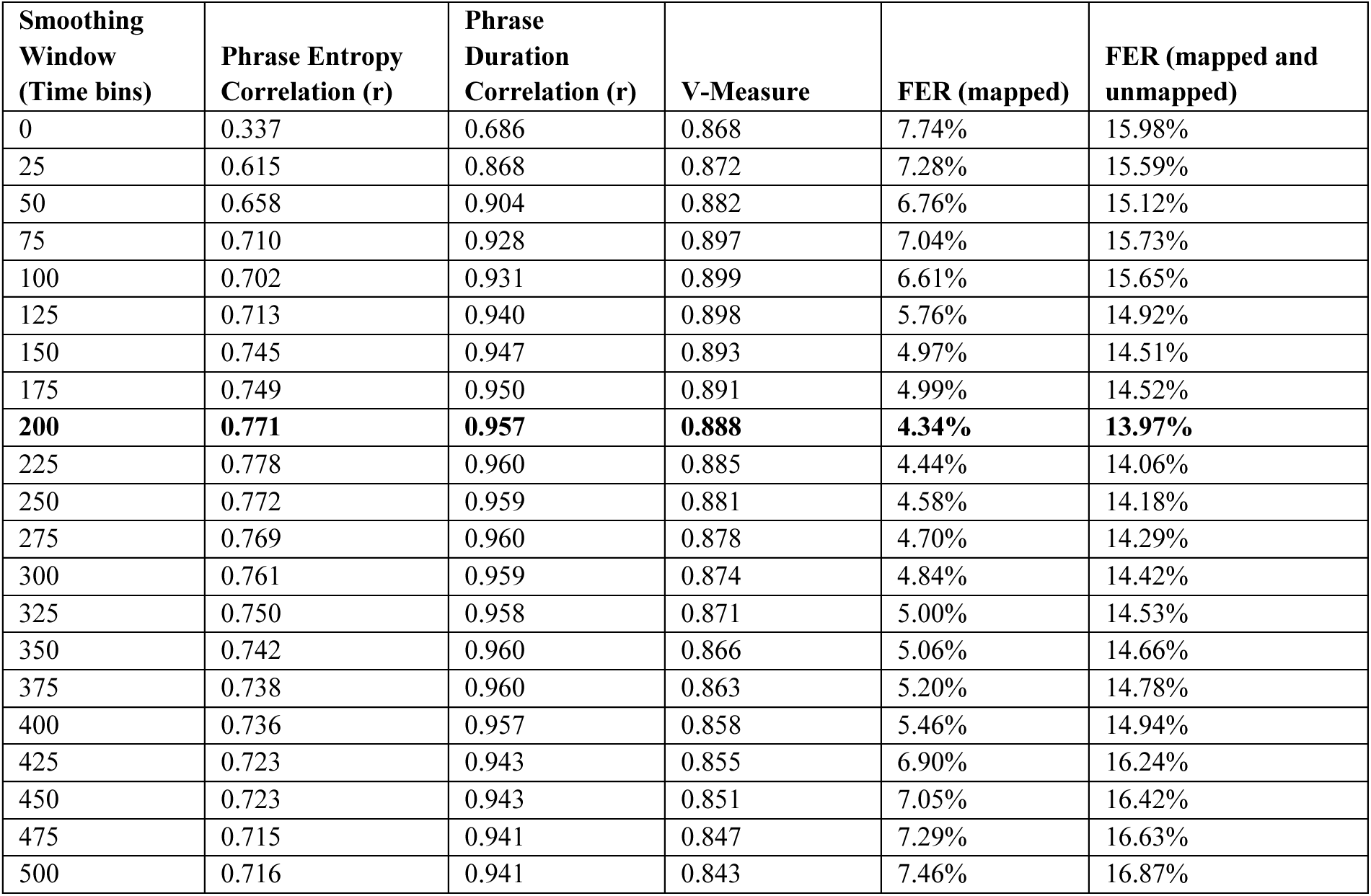
Raw data evaluating the impact of smoothing window size on clustering performance. Pearson correlations between HDBSCAN-derived and ground truth measures of phrase entropy and phrase duration, V-measure scores, and Frame Error Rates.

| **Smoothing Window (Time bins)** | **Phrase Entropy Correlation (r)** | **Phrase Duration Correlation (r)** | **V-Measure** | **FER (mapped)** | **FER (mapped and unmapped)** |
| --- | --- | --- | --- | --- | --- |
| 0 | 0.337 | 0.686 | 0.868 | 7.74% | 15.98% |
| 25 | 0.615 | 0.868 | 0.872 | 7.28% | 15.59% |
| 50 | 0.658 | 0.904 | 0.882 | 6.76% | 15.12% |
| 75 | 0.710 | 0.928 | 0.897 | 7.04% | 15.73% |
| 100 | 0.702 | 0.931 | 0.899 | 6.61% | 15.65% |
| 125 | 0.713 | 0.940 | 0.898 | 5.76% | 14.92% |
| 150 | 0.745 | 0.947 | 0.893 | 4.97% | 14.51% |
| 175 | 0.749 | 0.950 | 0.891 | 4.99% | 14.52% |
| **200** | **0.771** | **0.957** | **0.888** | **4.34%** | **13.97%** |
| 225 | 0.778 | 0.960 | 0.885 | 4.44% | 14.06% |
| 250 | 0.772 | 0.959 | 0.881 | 4.58% | 14.18% |
| 275 | 0.769 | 0.960 | 0.878 | 4.70% | 14.29% |
| 300 | 0.761 | 0.959 | 0.874 | 4.84% | 14.42% |
| 325 | 0.750 | 0.958 | 0.871 | 5.00% | 14.53% |
| 350 | 0.742 | 0.960 | 0.866 | 5.06% | 14.66% |
| 375 | 0.738 | 0.960 | 0.863 | 5.20% | 14.78% |
| 400 | 0.736 | 0.957 | 0.858 | 5.46% | 14.94% |
| 425 | 0.723 | 0.943 | 0.855 | 6.90% | 16.24% |
| 450 | 0.723 | 0.943 | 0.851 | 7.05% | 16.42% |
| 475 | 0.715 | 0.941 | 0.847 | 7.29% | 16.63% |
| 500 | 0.716 | 0.941 | 0.843 | 7.46% | 16.87% |

**Supplementary Fig. 6.**
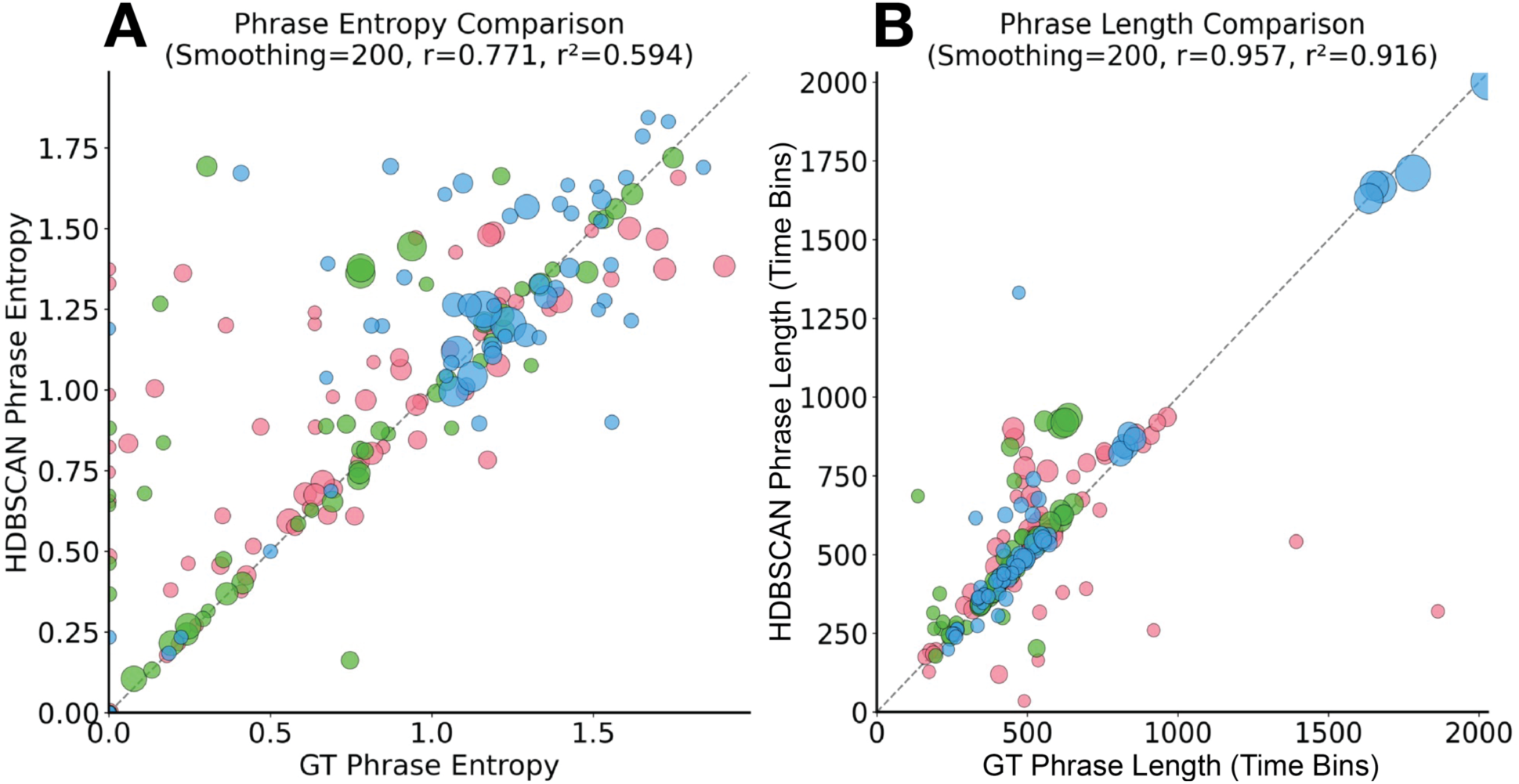
Phrase entropy correlations for each individual phrase label across three birds. Relationship between TweetyBERT-derived phrase metrics and human-derived ground truth (GT) metrics for three birds (colors represent individual birds). **(A)** Phrase entropy correlations **(B)** Phrase duration correlations. All analysis shown here were conducted with a 200 time bin smoothing window.

**Supplementary Table 7.** V-measure scores for layer and sublayer embeddings of TweetyBERT evaluated using UMAP projections from three birds in the TweetyNET dataset. Scores represent averages across these test birds. (A) V-measure scores for each layer-sublayer combination. The selected representation (Layer 3 Attention Output) ranks fourth among all combinations; however, the difference from higher-ranked combinations is minimal and likely within measurement noise. (B) Crucially, Attention Output is the highest-performing sublayer averaged across all layers, and (C) Layer 3 provides the strongest overall representations when averaged across all sublayers, supporting its selection for downstream analyses.

| Layer | Sublayer | V-Measure | Dim. |
| --- | --- | --- | --- |
| 3 | Feed Forward Output | 0.8935 | 196 |
| 4 | Intermediate Residual Stream | 0.8890 | 196 |
| 3 | Intermediate Residual Stream | 0.8859 | 196 |
| 3 | Attention Output | 0.8843 | 196 |
| 3 | Feed Forward Output GELU | 0.8829 | 768 |
| 2 | Feed Forward Output | 0.8757 | 196 |
| 2 | Attention Output | 0.8729 | 196 |
| 2 | Feed Forward Output GELU | 0.8619 | 768 |
| 1 | Attention Output | 0.8557 | 196 |
| 4 | Attention Output | 0.8494 | 196 |
| 4 | Feed Forward Output | 0.8420 | 196 |
| 2 | Intermediate Residual Stream | 0.8405 | 196 |
| 1 | Feed Forward Output GELU | 0.8400 | 768 |
| 1 | Feed Forward Output | 0.8326 | 196 |
| 4 | Feed Forward Output GELU | 0.8136 | 768 |
| 1 | Intermediate Residual Stream | 0.7518 | 196 |

| Sublayer | Avg. V-Measure | Dim. |
| --- | --- | --- |
| Attention Output | 0.8656 | 196 |
| Feed Forward Output | 0.8609 | 196 |
| Feed Forward Output GELU | 0.8496 | 768 |
| Intermediate Residual Stream | 0.8418 | 196 |

| Layer | Avg. V-Measure |
| --- | --- |
| 3 | 0.8866 |
| 2 | 0.8627 |
| 4 | 0.8485 |
| 1 | 0.8201 |

